# CompoundRay: An open-source tool for high-speed and high-fidelity rendering of compound eyes

**DOI:** 10.1101/2021.09.20.461066

**Authors:** Blayze Millward, Steve Maddock, Michael Mangan

**Affiliations:** Dept of Computer Science, The University of Sheffield, Sheffield, UK

## Abstract

Revealing the functioning of non-standard visual systems such as compound eyes is of interest to biologists and engineers alike. A key investigative method is to replicate the sensory apparatus using artificial systems, allowing for investigation of the visual information that drives animal behaviour when exposed to environmental cues. To date, ‘Compound Eye Models’ (CEMs) have largely explored the impact of features such as spectral sensitivity, field of view, and angular resolution on behaviour. Yet, the role of shape and overall structure have been largely overlooked due to modelling complexity. However, modern real-time raytracing technologies are enabling the construction of a new generation of computationally fast, high-fidelity CEMs. This work introduces new open-source CEM software (*CompoundRay*) alongside standardised usage techniques, while also discussing the difficulties inherent with visual data display and analysis of compound eye perceptual data. *CompoundRay* is capable of accurately rendering the visual perspective of a desert ant at over 5,000 frames per second in a 3D mapped natural environment. It supports ommatidial arrangements at arbitrary positions with per-ommatidial heterogeneity.

## Introduction

Insects visually solve an array of complex problems including the detection and tracking of fast moving prey *Wiederman et al*. (*2017*), long distance navigation *Wehner* (*2020*) and even 3D depth estimation *Nityananda et al*. (*2018*). These capabilities are realised using sensory apparatus that is fundamentally different from those of mammals. Therefore revealing the functional properties of the insect visual system offers insights for biologists as well as inspiration for engineers looking to develop novel artificial imaging systems *Land and Fernald* (*1992*); *Land* (*1997*); *Arendt* (*2003*); *Song et al*. (*2013*).

Arthropods possess two primary visual sensors known as compound eyes. Each eye is constructed from a patchwork of self-contained light-sensing structures known as ommatidia, each featuring a lens, a light guide and a cluster of photosensitive cells (Figure 1a). Ommatidia are physically interlocked with their neighbours, together forming a bulbous outer structure (the compound eye itself) that can vary in size, shape, and curvature, offering a range of adaptations for particular tasks and environments *Land and Nilsson* (*2002*) (Figure 1b). The properties of the lens and photo-receptors are fixed for individual ommatidia but can vary across regions of an individual’s eye *Meyer and Labhart* (*1993*) as well as between individuals of different castes *Collett and Land* (*1975*) and species *Land* (*1989*). This arrangement of independent, interlocked light sensing elements and lenses differs greatly from the mammalian system that utilises a single lens to project a high-resolution image onto a retina.

**Figure 1.**
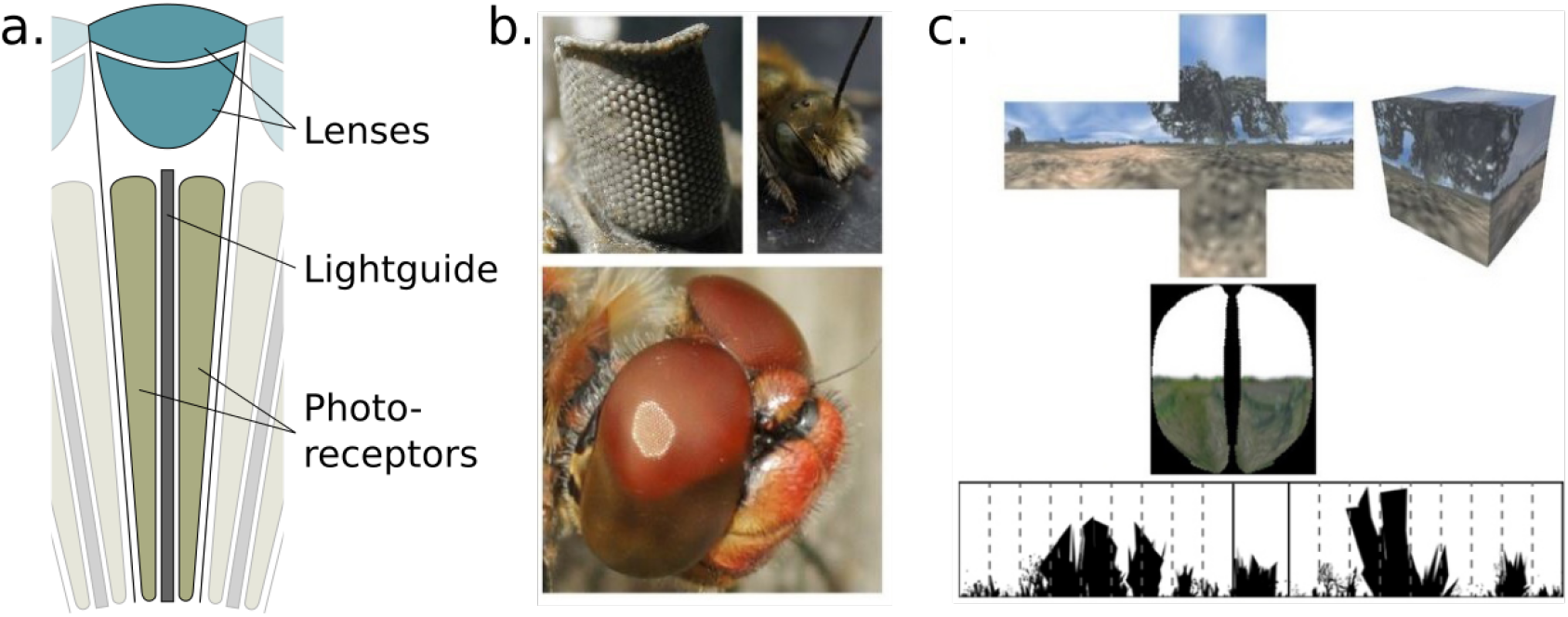
**a.** A diagram of a single ommatidium, seen with lensing apparatus at the top that guides light into the photo-sensitive cells below **b.** Images of real compound eyes consisting of hundreds to thousands of ommatidia. Upper right and lower image credits *Matthew Barber*. Upper Left image credit *Moussa Direct Ltd.* under the CC BY-SA 3.0 license. **c.** Previous compound eye-view recreation methods (*Neumann* (*2002*); *Polster et al*. (*2018*); *Wystrach et al*. (*2016*), top to bottom).

Creation of compound eye models (CEMs), in both hardware or software, are a well-established mechanism to explore the information provided by compound eyes and assess their impact on behaviour. Insights derived from this methodology include demonstration of the benefits of a wide field of view and low resolution for visual navigation *Zeil et al*. (*2003*); *Vardy and Moller* (*2005*); *Mangan and Webb* (*2009*); *Wystrach et al*. (*2016*), and the role played by non-visible light sensing for place recognition *MÖLLER* (*2002*); *Stone et al*. (*2014*); *Differt and Möller* (*2016*) and direction sensing *Gkanias et al*. (*2019*); *Lambrinos et al*. (*1997*). Yet, simulated CEMs tend to suffer from a common design shortcoming that limits their ability to accurately replicate insect vision. Specifically, despite differences in the sampling techniques used (e.g. see *Baddeley et al*. (*2012*); *Mangan and Webb* (*2009*) for custom sampling approaches, *Neumann* (*2002*); *Basten and Mallot* (*2010*) for rendering based cubemapping CEMs, and *Polster et al*. (*2018*) for ray-casting methods) all contemporary CEMs sample from a single nodal point. In contrast, the distributed arrangement of ommatidia on distinct 3D eye surfaces provides insects with a multi-nodal system that generates different information for different eye shapes.

To facilitate exploration of such features, an ideal rendering system would allow light to be sampled from different 3D locations through individually configured ommitidia replicating the complex structure of real compound eyes. High-fidelity compound-vision rendering engines with some of these features (though notably slower than real-time) were developed previously *Giger* (*1996*); *Collins* (*1998*); *Polster et al*. (*2018*), but were not widely adopted. Such issues are rapidly diminishing as dedicated ray-casting hardware emerges (e.g. Nvidia (Santa Clara, California, United States) *RTX* series GPUs *Purcell et al*. (*2005*); *Burgess* (*2020*)) that allows for the capture of visual data from multiple locations in parallel at high speed. Such systems present an ideal tool to replicate insect vision in unprecedented accuracy fast enough to allow effective exploration of the compound eye design space itself *Millward et al*. (*2020*). A next-generation insect eye renderer should:

1. Allow for the arrangement of an arbitrary number of ommatidia at arbitrary 3D points.
2. Allow for the configuration of individual ommatidial properties (such as lensing arrangement or spectral sensitivity).
3. Perform beyond real-time to allow exploration of the design space.

This paper presents a ray-casting based renderer, *CompoundRay*, that leverages modern hardware-accelerated ray-tracing graphics pipelines to capture all three of these criteria, allowing researchers in the fields of compound vision and biorobotics to quickly explore the impact varying eye designs have on an autonomous agent’s ability to perceive the world.

## Materials and Methods

### Raycasting-based Insect Eye Renderer

The majority of real-time rendering approaches are projection-based approaches inspired by the simple *pin-hole* camera. As Figure 2b shows, their images are derived by directly projecting the faces and vertices of the scene’s 3D geometry through a singular focal point and on to a projection plane. However, this approach (*polygon projection*) lacks the ability to simulate real-world light transport (and thus, lensing systems) accurately – in particular the spreading of light through reflection, refraction and diffraction, all of which have to be approximated using resource-heavy workarounds such as shadow maps *Stamminger and Drettakis* (*2002*), reflection maps *Miller and Hoffman* (*1984*) or other lookup maps for large precomputed data *Lopes and Fernandes* (*2014*); *Schäfer et al*. (*2012*).

**Figure 2.**
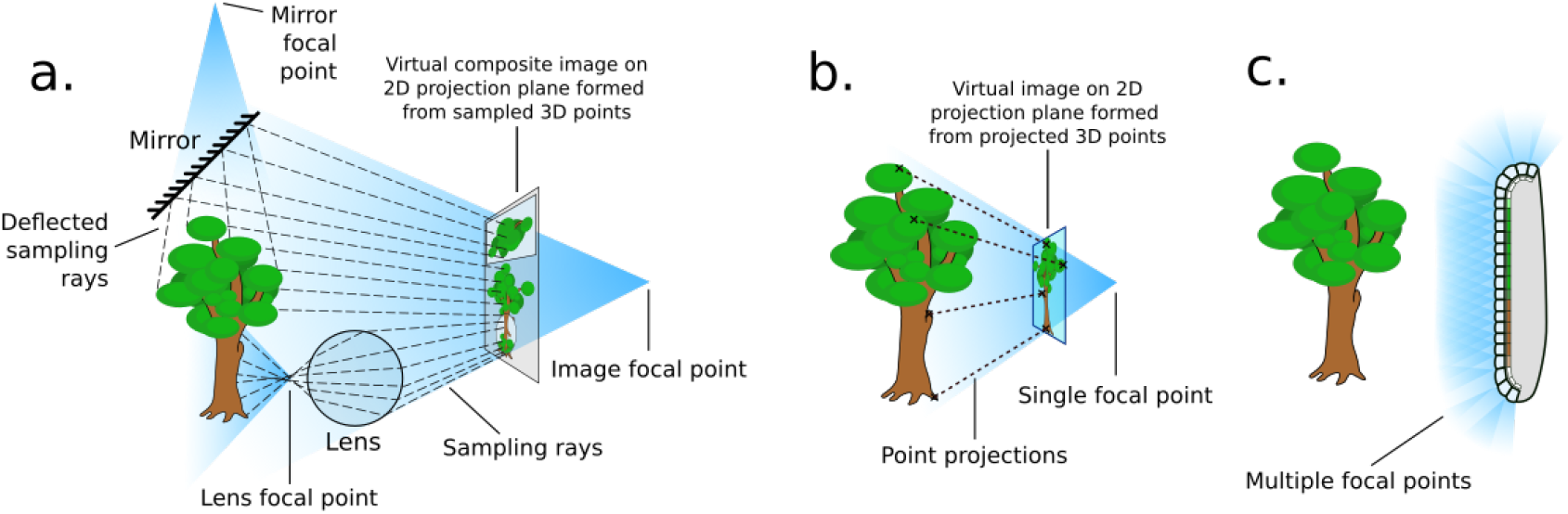
How differing rendering approaches form differing focal points. **a.** In 3D raycasting-based rendering rays are emmitted into a scene and *sample* the 3D geometry they contact, potentially forming many focal points. **b.** In 3D projection-based rendering, the polygons that comprise the 3D geometry are projected down to a viewing plane, through one focal point. **c.** A compound eye requires many focal points in order to image its surroundings.

In the real world, practically all light has been subject to at least one of these properties – for instance, objects in shadow are still visible due to interreflections *Langer* (*1999*), and glass or heat gradients can act as refracting lenses *Primmerman et al*. (*1991*). A polygon projection approach will struggle with these optical phenomena as the process directly transforms the faces of the 3D object from the scene onto the camera’s view plane, in effect applying a projection transform onto a plane or other regular surface forming a singular focal point (Figure 2b). In contrast to this, lenses or reflective surfaces in a real scene effectively act to create additional, sometimes complex, projection surfaces with multiple focal points that combine to form the final composite-perspective view seen by an observer (Figure 2a).

Ray-based methods offer an alternative to projective transform rendering: rays are sent out from the virtual view-point, simulating – in reverse – the paths of photons from the scene into the cone of vision, accumulating colour from the surface they interact with (Figure 2a). This allows for surfaces such as mirrors and lenses to accurately reflect light being rendered. The term *ray-based methods* here is used as an umbrella term for all rendering approaches that primarily use the intersection of rays and a scene in order to generate an image. In particular, we refer to *ray casting* as the act of sampling a scene using a single ray (as per *Roth* (*1982*)) – a process that CompoundRay performs multiple times from each ommatidium – and *ray tracing* as the act of using multiple recursive ray casts to simulate light bouncing around a scene and off of objects within, similar to early work in the field of shading *Appel* (*1968*); *Whitted et al*. (*1979*).

These approaches are incredibly computationally complex, as each ray (of which there can be thousands or even millions per pixel) needs to be tested against every object in the scene (which can itself be composed of millions of objects) to see if they intersect. Furthermore, any reflective objects require further intersections as they reflect light rays back into the scene. Historically, these ray-based methods have been limited to use in offline cases where individual frames of an animation can take many hours to render, such as in cinema *Christensen et al*. (*2018*), and prior to that in architectural and design drafting *Appel* (*1968*); *Roth* (*1982*). In recent years, however, graphics processing units (GPUs) have been increasing in capability. In order to better capture the photorealistic offerings of ray-based rendering methods, GPU manufacturers have introduced dedicated programmable ray-tracing hardware into their graphics pipelines *Purcell et al*. (*2005*). These so-called “RT cores” allow for efficient parallel triangle-line intersection to be performed, allowing for billions of rays to be cast – in real-time – into scenes even as complex as those found in modern game environments.

As compound eyes consist of a complex lensing structure formed over a non-uniform surface, they naturally form a multitude of focal points across a surface (Figure 2c). These projection surfaces are often unique and varied, meaning it is practically infeasible to find a single projective transform to produce the appropriate composite view in a single projection-based operation. As a result, ray-based methods become the natural choice for simulating compound vision, as opposed to the more commonly seen projection-based methods. By utilising modern hardware it is possible to effectively capture the optical features of the compound eye from the ommatidia up, in a high-fidelity yet performant way.

#### Modelling Individual Ommnitidia

As the compound eyes of insects contain many hundreds or even thousands of lenses (Figure 1b), and each acts to focus light to its own point (forming a unique perspective), ray-based methods become the natural technology to use to implement a simulator that can capture their unique sensory abilities. Here, a single ommatidia is first examined, simulated and then integrated into a complete compound eye simulation.

Each individual ommatidium in a compound eye captures light from a cone of vision (defined by the *ommatidial acceptance angle*), much in the way that our own eyes observe only a forward cone of the world around us. However, in the case of the ommatidium all light captured within the cone of vision is focused to a singular point on a small photoreceptor cluster, rather than the many millions of photoreceptors in the human eye that allow for the high-resolution image that we experience – in this way, the vision of a singular ommatidium is more akin to the singular averaged colour of all that is visible within its given cone of vision – it’s *sampling domain*, much as a pixel from a photo is the average colour of all that lies ‘behind’ it.

All objects within the ommatidium’s sampling domain will invariable impart influence on the final amount of sensed light, the total of which must be integrated, in much the same way that the *rendering equation* *Kajiya* (*1986*); *Immel et al*. (*1986*) is in traditional computer graphics – using a Monte Carlo approximation *Kajiya* (*1986*). Within a computer it is practically impossible to account for all infinite light rays converging on to a point, as such the amount of light received across the angular space of the viewing cone (which must be accrued in alignment with the the ommatidium’s *sampling function* – as the lens gathers more light from forward angles than those diverging away from the ommatidial axis) can be *sampled* a discrete number of times to approximate the entire sampling domain. Increasing the sample count will result in a more accurate estimate of the light converging on the point.

Previous works *Collins* (*1998*); *Polster et al*. (*2018*) have implemented this approximation of the light sampling function by assuming the influence of individual light rays, distributed statically across the sampling cone, is modulated dependent on the angle to the ommatidial axis via a Gaussian function with a full width at half maximum (FWHM) equal to the acceptance angle of the ommatidium, as seen in Figure 3a. *CompoundRay* also uses the Gaussian function as a basis for sampling modulation, however, unlike other works it is not approximated via a static sampling pattern with weighting biasing the forward direction (as is seen in 3a:i). Instead, sampling *direction* is modulated and chosen at random in relation to the Gaussian probability curve of the sampling function on a temporal, frame-by-frame basis. Each sample is assumed to have an equal weight, with the visual input to a single ommatidium then being formed as the average of all samples (Figure 3a:ii).

**Figure 3.**
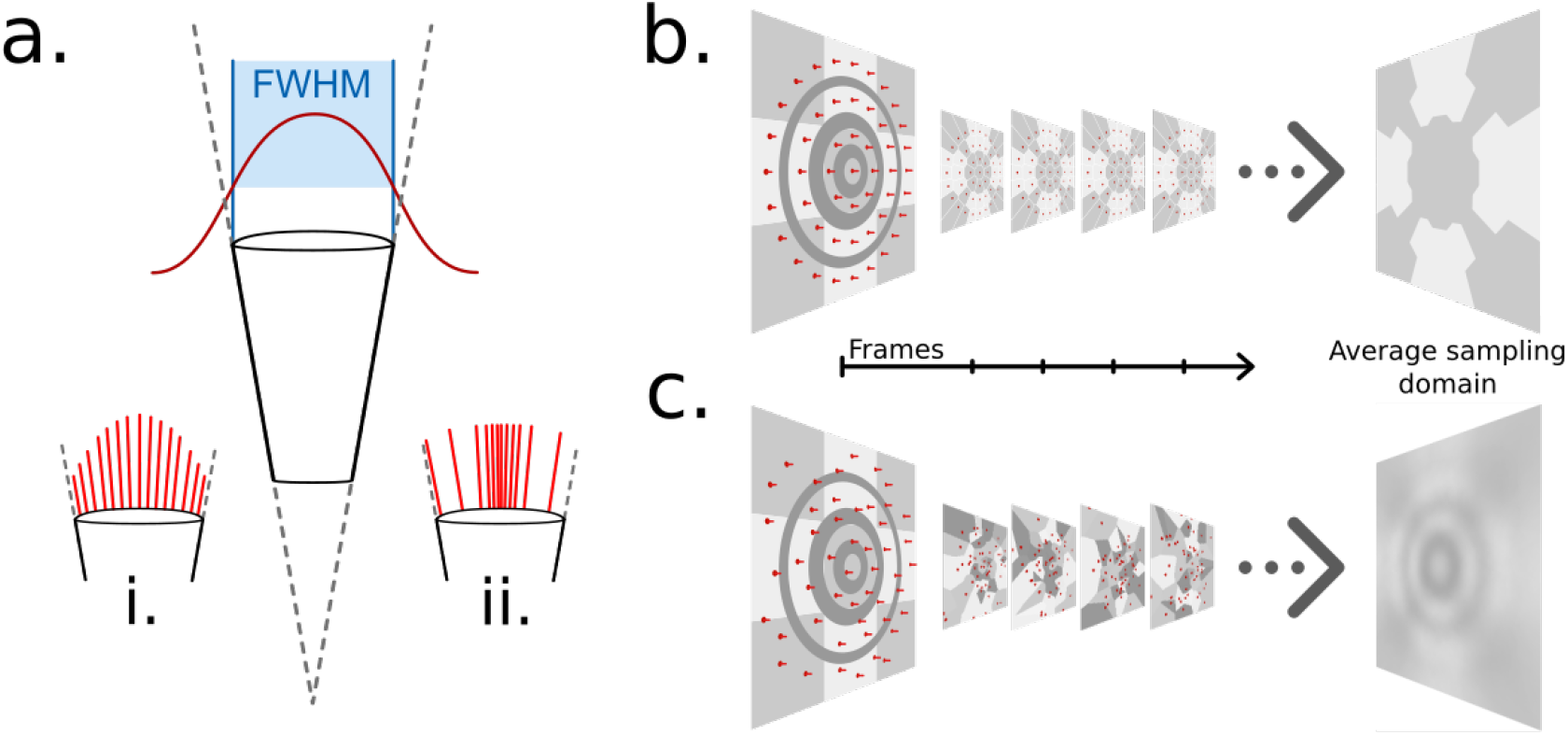
**a.** The gaussian sampling curve (red) with a full width at half maximum (FWHM) equal to the acceptance angle of each ommatidium. **i.** Example statically distributed sampling rays, length indicates weight. **ii.** Example stochastically distributed sampling rays, equal weight, distributed according to guassian distribution. **b.** A concentric circular static sampling pattern (red dots) aliases away the 3 concentric dark-grey rings present on a surface. **c.** The per-sample colours as seen from a single ommatidium mapping to the samples in *b*, note the lack of any influence from the dark-grey concentric rings, which have been aliased out.

Temporal modulation eliminates the potential of structured aliasing forming, as demonstrated in Figures 3b&c whereby, for instance in the case where per-ommatidium sampling is performed using static concentric circles, circular patterns matching the frequency would be aliased out of the ommatidium’s sampling domain if occurring out-of-phase with the sampling points. Increasing per-ommatidium sampling rate *can* ease these aliasing problems (even in the case of a static sampling pattern) at close range, but only serves to delay triggering structured aliasing until further distances where structural frequency matches sampling pattern again.

Choosing to stochastically vary the sampled direction on a frame-by-frame basis removes the structured nature of any single frame’s aliasing, as that same view point will generate different aliasing artifacts on the next frame, having the net effect of reducing structured aliasing when considering sequences of frames (Figure 3c) rather than single frames at a time, as opposed to any static sampling method (Figure 3b). One benefit of this approach is that averaging renderings from the same perspective becomes synonymous with an increased sampling rate, indeed, this is how the renderer increases sampling rate internally: by increasing the rendering volume for the compound perspective, effectively batch-rendering and averaging frames from the same point in time. However, as each ommatidium will sample the simulated environment differently for every frame rendered, small variations in the colour received at any given ommatidium occur over time. This point is explored further in the results section.

In the natural compound eye, light focused from each ommatidium’s sampling domain is transported via a crystalline cone and one or more rhabdomeres toward the receiving photoreceptor cluster – a process that has been modelled in past works *Song et al*. (*2009*). However, modelling the biophysical intricacies of the internal ommatidium structure is considered beyond the scope of this software, and is considered a post-processing step that could be run on the generated image, considering each pixel intensity as a likelihood measure of photon arrival at an ommatidium.

#### From single ommatidia to full compound eye

By arranging the simulated ommatidia within 3D space so as to mimic the way real compound eyes are arranged, an image can be generated that captures the full compound eye view. For this task, per-ommatidium facet diameter and acceptance angle can be used to generate the cone of influence of each ommatidium, and simulated ommatidia can be placed at any position and orientation within the environment. As the system is based on ray-tracing techniques, orientation and position can be set with a single per-ommatidium data point, effectively allowing each ommatidium to act independently of each other. This has the benefit of allowing the system to simulate any type of eye surface shape, and even multiple eyes simultaneously, fulfilling the first of the three criteria: *Allowing for the arrangement of an arbitrary number of ommatidia at arbitrary 3D points*.

### The CompoundRay Software Pipeline

The renderer is written in C++ and the Nvidia CUDA GPU programming language and allows for a 3D environment representing a given experimental setup to be rendered from the perspective of an insect. The core of the renderer runs in parallel on the GPU (the *device*) as a series of CUDA shader programs, which are driven by a C++ program running on the host computer (the *host*). Environments are stored in *GL Transmission Format* (glTF) *Robinet et al*. (*2014*) files consisting of 3D objects and cameras, the latter of which can be of two types: compound (structured sets of ommatidia) or traditional (perspective, orthographic and panoramic). Traditional cameras are implemented to aid the user in the design and analysis of their assembled 3D environment. Compound cameras contain all relevant information for rendering a view of the environment from the perspective of a compound eye. Each camera stores the information required for rendering its view (such as its orientation, position, field of view or ommatidial structure) in an on-device **data record** data structure.

Figure 4 shows the operational design of the renderer from the device side. The renderer is built on top of Nvidia’s *Optix* *Parker et al*. (*2010*) raytracing framework, which is a **pipeline**-based system: a pipeline is assembled, which then allows for parallel per-pixel rendering of the environment. A **pipeline** consists of a **ray generator shader**, a **geometry acceleration structure (GAS)** and numerous **material shaders**. A shader is a small program that is designed to be run in a massively-parallel fashion. In a typical application, an individual shader program will be loaded onto a GPU and many thousands of instances of the program will be executed, with varying input parameters. The returned values of each instance are then returned to the ‘host-side’ of the application, allowing for the optimisation of tasks that are well-suited to parallelisation.

**Figure 4.**
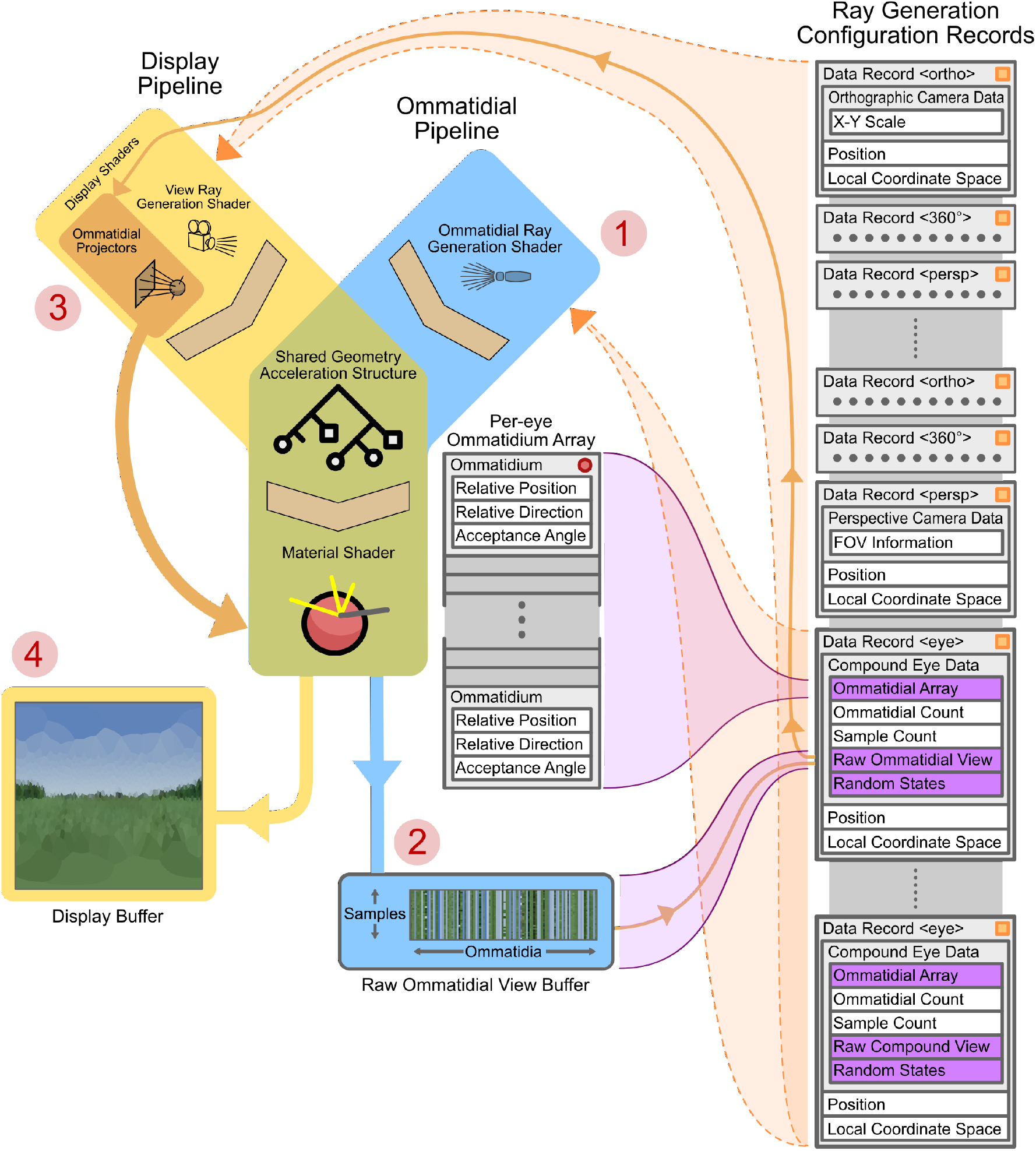
GPU-side renderer operational diagram. Featured are the joined *ommatidial* and *display* rendering pipelines (including display buffers) as well as the on-device per-camera configuration records and related memory allocations. Data-structure memory allocations are shown in grey (indicating in-place data) or purple (indicating pointers to other sections of data). Vertical stacks of memory indicate contiguously allocated structures. Data-structures marked with an orange square are automatically updated as their host-side copies change; those with red circles are manually updated via API calls. Circled numbers 1 to 4 indicate the path of processing through both pipelines required to display a compound eye view.

A **ray generator shader** spawns rays to sample the environment with – these are dependent on the type, position and orientation of the chosen camera. In the case of a panoramic camera (in the display pipeline), for instance, rays are spawned in a spherical manner around a central point (the *position* component of the currently selected camera’s ray generation configuration record). In the case of a compound eye (in the ommatidial pipeline), each ray is generated as a function of the relative position and direction of a single ommatidium to the eye’s position and orientation (note that the “eye”-type data records in Figure 4 contain links to per-eye ommatidial arrays that define the ommatidial configuration of the eye).

A **GAS** stores the 3D geometry of the scene on the device in a format optimised for ray-geometry intersection, and also provides functions to perform these intersection tests. Upon the intersection of a generated ray and the environment the material shader associated with the intersected geometry is run. These **material shaders** are responsible for the calculation and retrieval of the correct colour for the given geometry. For example, if a ray were to fall onto a blade of grass, the material shader would be expected to compute the green colour that could be seen, taking into account any light that might be affecting the scene. Currently CompoundRay implements only a simple texture lookup, interpolating the nearest pixels of the texture associated with the geometry intersected.

Two pipelines are used in the renderer: an **ommatidial pipeline** and a **display pipeline**. The **ommatidial pipeline** handles sampling of the environment through the currently active compound eye, saving all samples from all ommatidia to a buffer stored within the eye’s “Raw Ommatidial View” portion of its data record. First it generates per-ommatidium rays (Figure 4–1) via instances of the ommatidial ray generation shader. It then uses these to sample the environment through the GAS and appropriate material shaders, finally storing each ommatidium’s view as a vector of single-pixel tri-colour samples in the raw ommatidial view buffer (Figure 4–2).

Conversely, the **display pipeline** handles the generation of the user-facing **display buffer** (Figure 4–4). In the case of generating a compound eye view display, *ommatidial projector* shaders are spawned (one per output pixel) and used to lookup (Figure 4–3) and average the appropriate values from the per-camera raw ommatidial view buffer. These values are then re-projected into a human-interpretable view – in the case of the image seen at Figure 4–4, using a spherical orientation-wise Voronoi diagram projected into 2D by mapping angular yaw and pitch to 2D (x,y) coordinates. However for practical applications, this re-projection can be to a simple vector of all ommatidial values, allowing an algorithm to examine the data on a per-ommatidium basis. In the case where the current camera is non-compound the display pipeline becomes the only active pipeline, handling the initiation of ray generation, GAS intersection and materials shading for any given camera view in a single pass. Both pipelines share a common geometry acceleration structure and material shaders so as to save device memory.

A full compound eye rendering (as indicated via numbered red circles in Figure 4) consists of first rendering the ommatidial view via the ommatidial pipeline to the current camera’s raw ommatidial view buffer using the associated eye data record. After this, the raw ommatidial view buffer is re-projected onto the display buffer as a human-interpretable view via the ommatidial projector shaders within the display pipeline. Alternately, rendering using a traditional camera skips steps 1 and 2 and instead simply renders to the display view by passing fully through the display pipeline using the data record associated with the selected camera.

#### Scene Composure and Use

Using the glTF file format for scene storage allows the experimental configuration of 3D models, surface textures and camera poses to be packed into a single file (although per-eye ommatidial configurations are stored separately as CSVs and linked to within the glTF file). The glTF format is a JSON-based human-readable file format for 3D scene storage that is readily supported by a number of popular 3D editors and has allowances for extensions within its specification. As such, all configurable parts of the scene that relate to CompouundRay are placed within each camera’s reserved “extras” property. The renderer is programmed to ingest these extra properties, expanding the scene definition beyond standard glTF without compromising the readability of scene files by other third-party 3D modelling suites.

While the software comes packaged with a stand-alone executable that can load and render compound-eye scenes, this is not the most powerful way of using the tool. When compiled, a shared object library is also generated that can then be called from more accessible languages, such as Python *Van Rossum and Drake* (*2009*). Through the use of this, Python’s *ctypes* system and a helper library that is bundled with the code, it can be used directly with Python and the Numpy *Harris et al*. (*2020*) mathematical framework. This allows for a significantly wider experimental scope through automated configuration – for instance, one can programmatically move an insect eye around a simulated world extracting views directly from the renderer to be used alongside neural modelling frameworks accessible under Python. This is the recommended way of using the rendering system due to the increased utility that it affords.

## Results

In this section two 3D environments are used to benchmark and examine the performance of *CompoundRay*. The first is a lab environment inspired by those used in insect cognitive experiments *Ofstad et al*. (*2011*) and the second is a 3D scan of a real-world natural environment covering an area of a number of square kilometers. The natural environment was constructed using structure-from-motion photogrammetry using photos captured from a drone over rural Harpenden, England (3D model subject to upcoming publication, available via contact of Dr. Joe Woodgate, Queen Mary University of London).

### Criterion 1: Arbitrary arrangements of Ommatidia

As stated in criterion 1 of the required feature list (defined in the introduction), the renderer must support arbitrary arrangements of ommatidia within 3D space. Figure 5 shows two different environments rendered using three differing eye designs, from a simple spherical model with spherical uniformly distributed ommatidia that more closely align with current compound eye rendering methods, to a hypothetical arbitrary arrangement similar to that of the now long-extinct *Erbenochile erbeni*. While Figures 5a&b are still possible to simulate using traditional projection-based methods due to their ommatidial axes converging onto one central spot, 5c demonstrates an eye with multiple unique focal points across an irregular surface.

**Figure 5.**
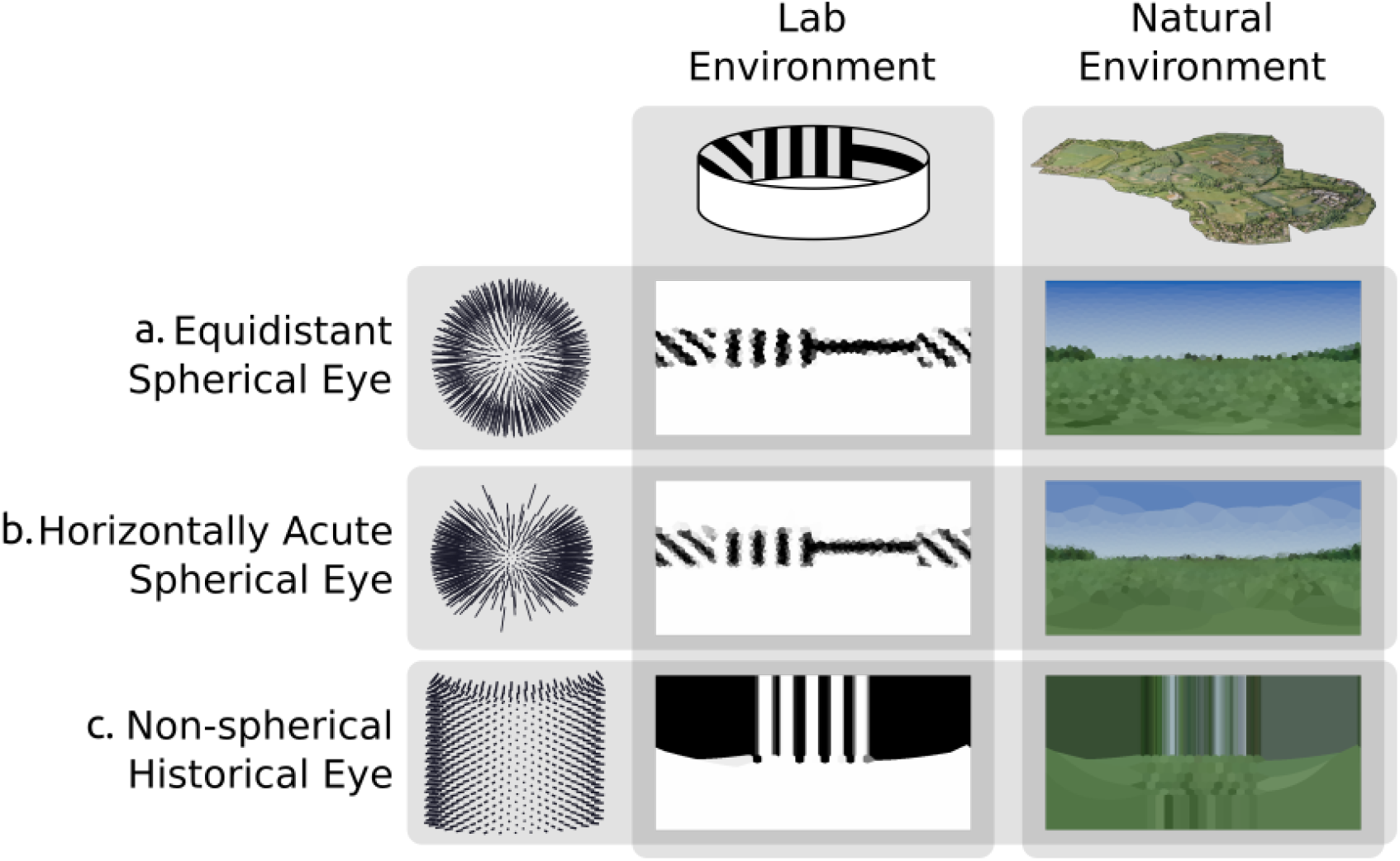
Rendering from eyes with ommatidia placed on arbitrary surfaces in one lab environment and one 3D scan of a natural outdoor environment. **a.** Spherically uniformly distributed ommatidia. **b.** Ommatidia clumped to form a horizontal acute zone **c.** A design inspired by the *Erbenochile erbeni*.

### Criterion 2: Inhomogeneous Ommnitidial Properties

Criterion 2 states that it should be possible to specify heterogeneous ommatidial optical properties across an eye. As demonstrated in Figure 6, the renderer achieves this by allowing for shaping of the acceptance cone of each ommatidium. In doing so, Figure 6 also demonstrates the importance of heterogeneous ommatidial configurations.

**Figure 6.**
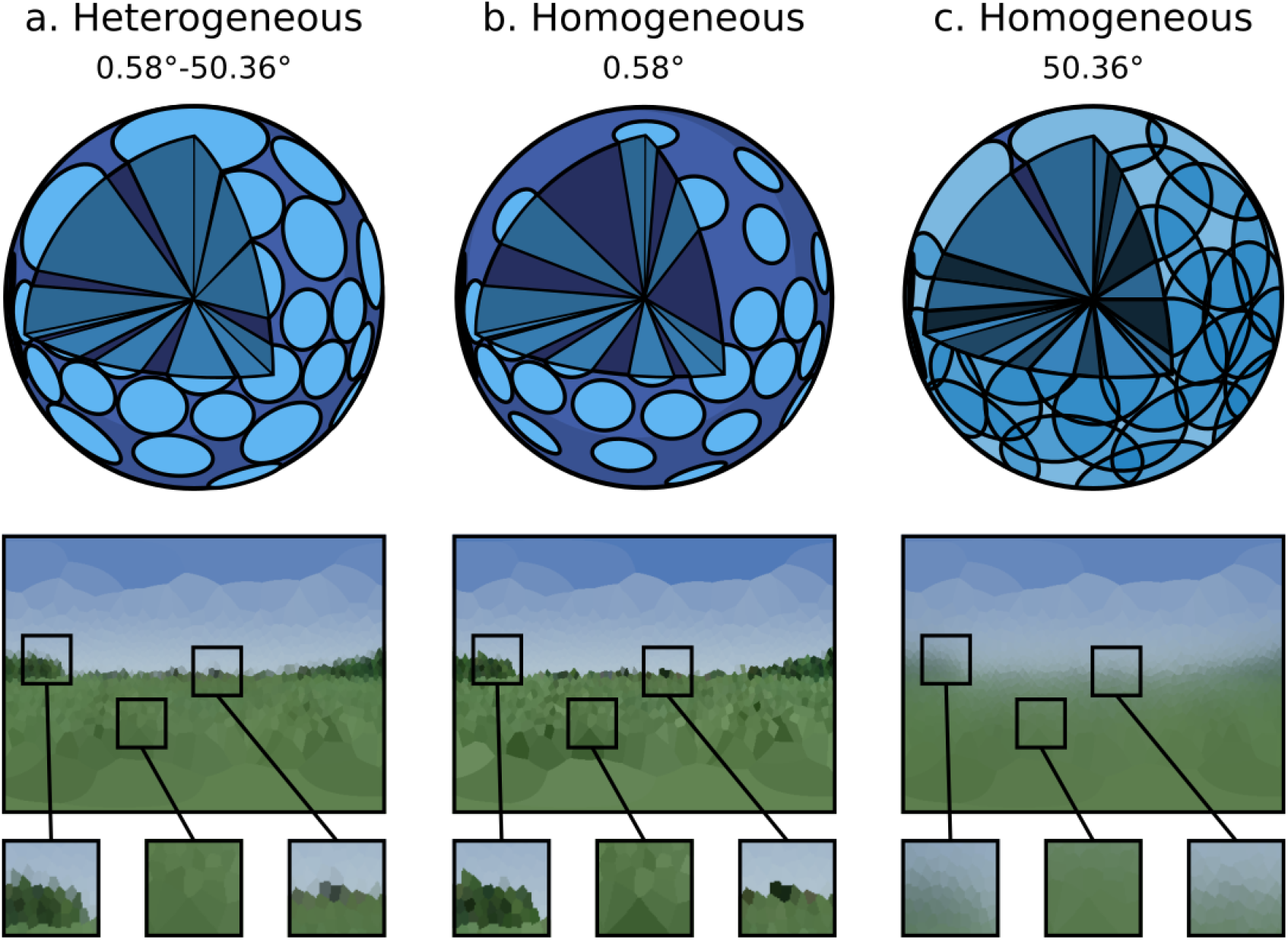
A single environment as viewed through: a. An eye with heterogeneous ommatidial acceptance angles. **b&c.** Eye designs with homogeneous ommatidial acceptance angles, note the sharp aliased edges in *b* and the blurred edges in *c*. Above each eye view is a diagram demonstrating the type of coverage allowed by each design into the eye’s spherical field of view, showing the under- and over-sampled nature of *b* & *c* compared to *a*.

Figures 6b&c show a non-uniform eye design with homogeneous, globally-identical, ommatidial acceptance angles. It can be seen that in b, where the acceptance angle is lower, aliasing occurs as objects are only partially observed through any given ommatidium, causing blind spots to form, introducing noise into the image. Conversely, in c, where the acceptance angle is large, the image becomes blurry, with each ommatidium over-sampling portions of its neighbour’s cone of vision. Sub-figure a, however, demonstrates a heterogeneous ommatidial distribution, in which the acceptance angles toward the dorsal and ventral regions of the eye are larger compared with the angles of those around the eye’s horizontal acute zone. This mirrors what is seen in nature, where ommatidial field of view is seen to vary in order to encompass the entirety of an insect’s spherical visual field *Land and Nilsson* (*2002*). As can be seen in the generated image, this is important as it appropriately avoids creating blind-spots in the dorsal and ventral regions while also avoiding over-sampling (blurring) in the horizontally acute region, resulting in a much clearer picture of the observed environment while also minimising the required number of ommatidia. This maximisation of utility while minimising structural complexity is a hallmark of compound eye design.

### Criterion 3: Speed

#### Minimum Sample Count

In order for rapid exploration and experimentation of the design space of compound eyes, criterion 3 (as defined in the introduction) requires that any such rendering system should be able to perform at real-time or faster. However, performance of the system is dependent on the number of rays required to render a given scene from a given eye; higher total ray counts will require more compute power. As discussed in the section *“Modelling Individual Ommatidia”*, per-ommatidial sampling choice can have a dramatic impact on the image generated, meaning that lower ray counts – while increasing performance – will decrease the accuracy of the image produced. As a result, it is important to be able to define some minimum number of samples required for any given eye in any given environment in a reliable and repeatable manner. Previous works *Polster et al*. (*2018*) have attempted to establish a baseline number of samples required per ommatidium before additional samples become redundant only via qualitative visual analysis of the results produced. Here we perform quantitative analysis of images produced by the renderer in order to more definitively define this baseline.

Unlike previous works, the temporally stochastic nature of *CompoundRay*s’ sampling approach can be used to aid the measurement of impact that sampling rate has on the final image. By measuring the per-ommatidium spread (here, standard deviation) of the Euclidean distance of the received colour as plotted in RGB colour space over a range of frames captured from a static compound camera, the precision of the light sampling function approximation can be measured. To control for the varying field of views (FOV) of each ommatidium (which impact the number of samples required, as larger FOVs require more sampling rays to accurately capture the ommatidium’s sampling domain), we normalize the measure of spread recorded at each ommatidium against its degree of coverage, in steradians.

Figure 7 shows this measure of spread within the compound eye across a range of samplerays-per-ommatidium, from 1 to 700. As the number of samples increases the spread decreases logarithmically. We propose iteratively increasing samples until the maximum spread is below some arbitrarily defined threshold value. Here, we use a threshold of 1%, meaning that the maximum expected standard deviation of the most deviant ommatidium on an eye (normalised to one steradian) should be no greater than 1% of the maximum difference between two received colours, here defined as the length of vector [255, 255, 255]. This method allows for standardised, eye-independent configuration of per-ommatidial sampling rate.

**Figure 7.**
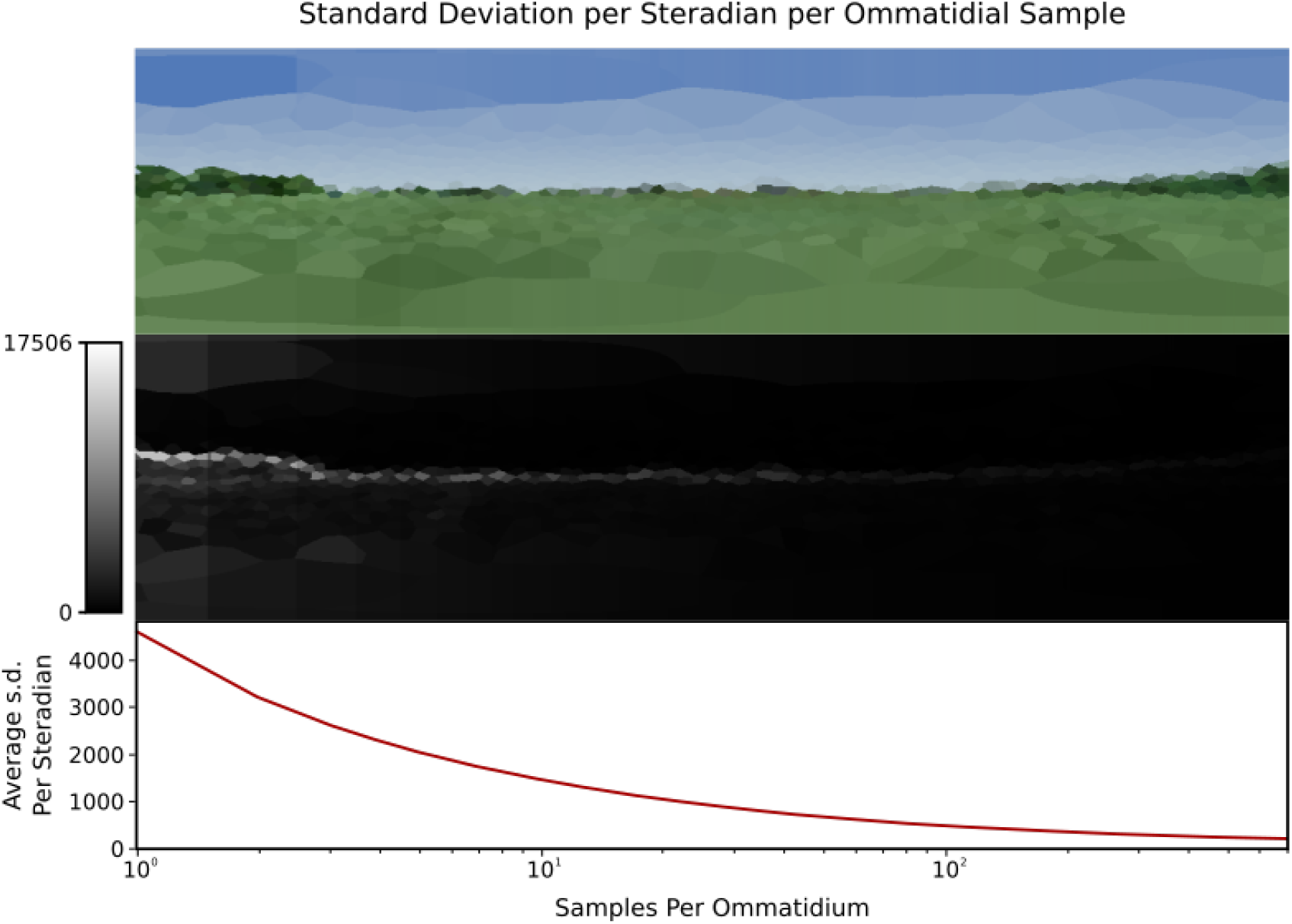
A single viewpoint viewed with 1 through to 700 samples per ommatidium, left to right, logarithmic scale. **Top**: how changing sample count changes the generated image. Note that the left side (with lower per-ommatidial samples) is more “jagged” due to aliasing. **Middle**: relative standard deviations of each ommatidium across 1000 frames, where difference is measured as the Euclidean distance between the two colours. **Bottom**: plot of the average standard deviation of an eye per sample rays per ommatidium, normalised to one steradian. Standard deviations decrease both on average (**bottom**) and per-ommatidia (**middle**) as per-ommatidium sampling rate increases.

However, the required number of samples per eye is highly positively correlated with the visual frequency and intensity of the simulated environment at any given point, with locations exhibiting high contrast seen from a distance requiring higher sampling rates in order to properly resolve. This can be seen in the the skyline in Figure 7, which remains dominant throughout the majority of the sampling rates, consistently forming the source of the highest spread. Figure 8 shows how the perceived variance changes over two example environments. The varying environment presents an obstacle in our attempts to standardise sampling rates, as deriving a suitable minimum sampling rate at one point in an environment – for example, where the visual frequency was, on average, low – could lead to aliasing occurring in more visually complex areas of the same environment, potentially introducing bias for or against certain areas if analysis on the generated data were sensitive to the variance introduced.

**Figure 8.**
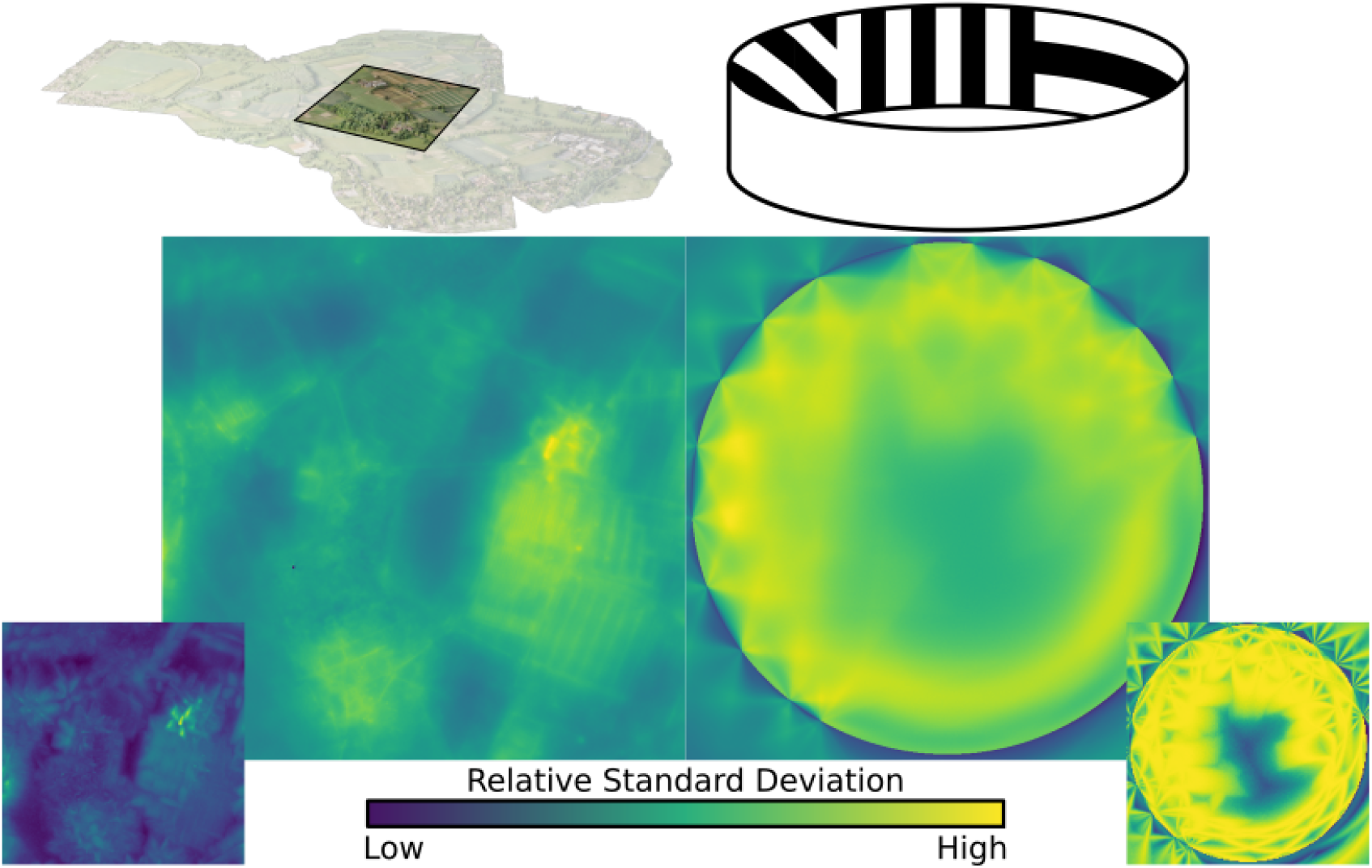
Standard deviation, averaged over all ommatidia in an eye, mapped over the natural and lab environments, showing a range of “hotspots” of high variance in renderings. These hotspots must be accounted for when calculating minimum sample ray count. **Inserts bottom left & right**: the maximum (over all ommatidia) standard deviation recorded in the eye at each location.

One method to attempt to control for this would be to search the environment for the location of highest variance and perform minimum sampling rate calculations there. This is the method this work follows, using a genetic algorithm to find the location of highest variance before iteratively increasing sampling rate until a frame-wise standard deviation of less than 1% is observed in the single ommatidium with the highest standard deviation. Alternatively, sampling rates could be updated in an online fashion by storing a rolling average of the maximum sample spread and increasing and decreasing sampling rate to keep the average below the considered acceptable spread percentage (here 1%).

#### Performance

Figure 9 shows the rendering speed, in frames-per-second, against the number of sample rays being emitted in each frame when running using a selection of high-end consumer-grade graphics cards and our two sample environments. Marked on the graph are the total per-frame sample counts required by common insect eyes, chosen to encourage a maximum per-ommatidium standard deviation of 1%.

**Figure 9.**
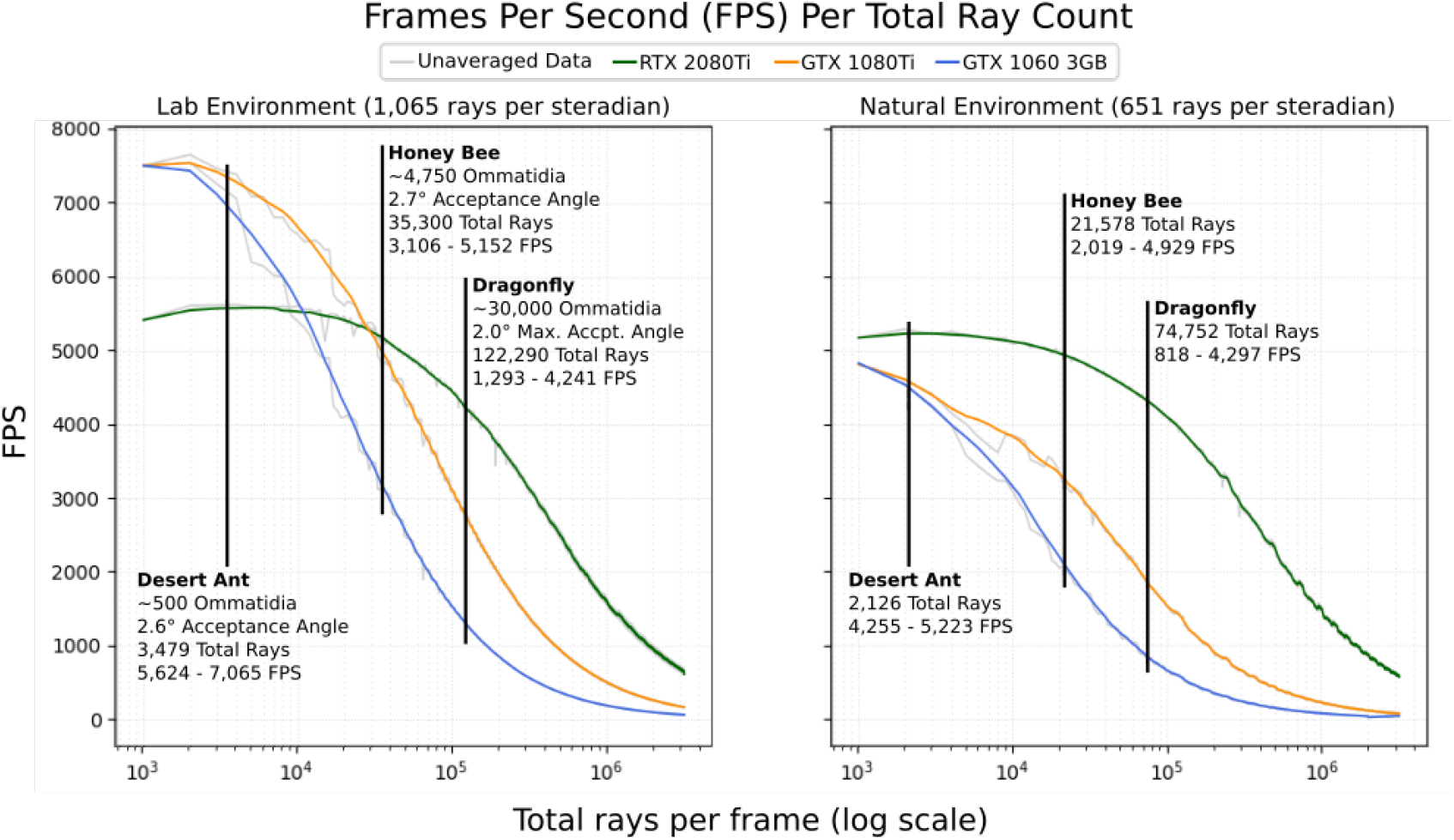
The average FPS (500 samples) per total number of rays dispatched into the scene for 3 different graphics cards in the two environments depicted in Figure 5. Marked are expected FPS values for a desert ant *Schwarz et al*. (*2011*), honey bee *Greiner et al*. (*2004*) and dragonfly *Labhart and Nilsson* (*1995*); *Land and Nilsson* (*2002*).

As can be seen, performance improves significantly when the renderer is able to utilise the next-generation ray-tracing hardware available in the newer RTX series of NVidia graphics cards, consistently out-performing older cards in the larger natural environment, and outperforming in the smaller lab environment when using more than 30,000 rays. Counter-intuitively, the previous generation of NVidia graphics cards were found to out-perform the RTX series in the (relatively small) lab environment at lower ray counts (below 30,000). This appears to be due to static over-head involved with the use of the RTX series’ ray-tracing cores, which may also be the cause of the small positive trend seen between 1,000 and 8,000 samples when using the 2080Ti. Another potentially counter-intuitive result that can be seen here is the higher performance of the dragonfly eye in the natural (high polygon) environment verses in the low-polygon laboratory environment. This is because the laboratory environment produces significantly higher visual contrast (emphasised in Figure 8) due to it’s stark black/white colour scheme, resulting in a requirement of almost double the number of rays per steradian to target 1% standard deviation, resulting in higher ray counts overall, particularly in models with a high number of ommatidia.

### Challenges of Visualisation and Eye Comparison

Compound eyes are inherently different from our own, and the CompoundRay rendering system allows for data collection on arbitrary compound surfaces. With this comes the risk of anthropomorphising the data received. At its base level, the eye data gathered is most ‘true’ when considered as a single vector of ommatidial samples, much in the same way that the neural systems of any insect will interpret the data, perhaps with some spatial relational information encoded. In contrast, all the images that we have presented here have been projected onto a 2D plane in order to be displayed. Great care must be taken when considering these projections, as they can introduce biases that may be easy to overlook. For instance, in Figures 5a&b, orientation-wise spherical projection mapping was used to artificially place the ommatidia across the 2D spaces, forming panoramic Voronoi plots of observed light at each ommatidium. In these projections, much as is seen in 2D projections of Earth, the points at the poles of the image are artificially inflated, giving a bias in terms of raw surface area affected toward ommatidia present in the ventral and dorsal regions of the eye.

In the case of the first two eye designs shown in Figure 5 this projection bias can be controlled for by using a projection method that controls for the area of a projected sphere into 2D space, such as the HEALPix *Górski et al*. (*2005*), Mollweide or other such equal-area projection schemes *Sousa et al*. (*2019*). However, this does not work for all eye designs. The design present in Figure 5c is inspired by that of the *Erbenochile erbeni* (1b, upper left), which exhibits a largely cylindrical shape. For this eye design, mapping onto a sphere (even using equal-area projection) would introduce bias, as either near-parallel ommatidia would occlude each another in the case of orientation-wise projection (as is seen happening in Figure 1c) or ommatidia further from the centre would be biased against as they are moved artificially closer together through projection onto the sphere than would happen to ommatidia on the same eye closer to the centre of the projection sphere.

These projection biases demonstrate how fundamentally difficult it is to compare eye designs that do not share an underlying surface structure, unless those surfaces happen to be affne transformations of one another (they are *affinely equivalent*). For any surfaces that are affnely equivalent, the surface itself can be used as a projection surface. In this way, for instance, the outputs of two cubic eyes could be compared to each other by performing all comparison calculations on the cube’s surface, interpolating between points where an ommatidia exists on one eye but not another. It is trivial to compare compound eye images from eyes with the same structure in terms of ommatidial placement, and eyes with surfaces that are affinely equivalent can also be compared, however further precautions must be taken when comparing the images from two or more eyes that do not have compatible projection surfaces so as not to unduly bias a portion of one surface over any other.

## Discussion

This paper has introduced a new compound eye perspective rendering software, *CompoundRay*, that addresses the need *Millward et al*. (*2020*); *Taylor and Baird* (*2017*); *Stürzl et al*. (*2015*) for a principled, fast, compound-perspective rendering pipeline. In particular, the tool supports arbitrary instantiation of heterogeneous ommatidia at any place in space, and can perform rendering tasks at framerates on the order of thousands per second using consumer-grade graphics hardware. It has demonstrated the utility afforded by the software and highlighted the importance of using higher-fidelity compound-eye perspective rendering systems by demonstrating the visual differences introduced by altering these often unaltered traits. It has also set a grounding for reasonable use of the software to ensure reproducibility and reduce biasing introduced as a result of the variety present in simulated environments and compound eye designs while warning of potential dangers that might arise during data analysis. With the introduction of the CompoundRay rendering library it is possible to simulate in a timely manner the visual experience of structurally diverse compound eyes in complex environments, digitally captured or otherwise manually constructed.

This work lays the technical foundations for future research into elements of compound eye design that have previously been given little consideration. In particular, CompoundRay can act as the previously missing step to integration of recently catalogued real eye designs *Baird and Taylor* (*2017*) and increasing capture of insect environments *Stürzl et al*. (*2015*); *Risse et al*. (*2018*) and routes *Risse et al*. (*2017*). Further to this, it can be used in the further exploration of recently emerging work surrounding microsaccadic sampling in drosophila *Juusola et al*. (*2017*), as well as aid with similar vision-centric neurological studies *Kemppainen et al*. (*2021*); *Wystrach et al*. (*2016*); *Viollet* (*2014*). On top of the potential biological insights CompoundRay may be able to offer, the ability to rapidly search the design space of compound eyes will enable faster iteration on insect-inspired physical sensors *Floreano et al*. (*2013*); *Song et al*. (*2013*), allowing for design experimentation before manufacture. The insights gained from more thoroughly exploring the insect visual perspective—and its design relative to visual feature extraction—will help guide the development of artificial visual systems by considering not only visual post-processing steps, but also the intrinsic structure and design of the image retrieval system itself.

The CompoundRay software is open-sourced on GitHub (at https://github.com/manganlab/eye-renderer) to allow for easy and systematic comparisons to be made between varying models of insect behaviour, without the introduction of implementation-specific differences in rendering of the insect perspective. In the future, the library can easily be extended to include further extensions to the modelling environment such as shadows, polarisation *Zeil et al*. (*2014*) or solar compass patterning in the virtual sky *Gkanias et al*. (*2019*).

## Acknowledgements

Thanks to Dr. James Knight (University of Sussex) for his input and feedback on the manuscript, and Dr. Joe Woodgate (Queen Mary University of London) for provision of the natural environment 3D model. Graphics cards were supplied by the Brains on Board research project (EP/P006094/1) and project partner NVidia. Grant funding was provided by EPSRC awards EP/P006094/1 and EP/S030964/1.

